# Effects of prefrontal tDCS on dopamine-mediated behavior and psychophysiology

**DOI:** 10.1101/697466

**Authors:** Michael J. Imburgio, Hannah K. Ballard, Astin C. Cornwall, Darrell A. Worthy, Jessica A. Bernard, Joseph M. Orr

**Author notes:** Corresponding author’s. Contributed Equally (Michael J. Imburgio, Hannah K. Ballard). Contributed Equally (Jessica A. Bernard, Joseph M. Orr).

## Abstract

The ability to manipulate dopamine *in vivo* through non-invasive, reversible mechanisms has the potential to impact clinical, translational, and basic research. Recent PET studies have demonstrated increased dopamine release in the striatum after prefrontal transcranial direct current stimulation (tDCS). We sought to extend this work by examining whether prefrontal tDCS could demonstrate an effect on behavioral and physiological correlates of subcortical dopamine activity. We conducted a between-subjects study (n = 30) with active and sham tDCS and used spontaneous eye blink rate (EBR), facial attractiveness ratings, and greyscales orienting bias as indirect proxies for dopamine functioning. The initial design and analyses were pre-registered (https://osf.io/gmnpc). Stimulation did not significantly affect any of the three measures, though effect sizes were often moderately large and were all in the predicted directions. Additional exploratory analyses suggested that stimulation’s effect on EBR might depend on pre-stimulation dopamine levels. Our results shed light on the sensitivity of indirect measures of dopamine in humans and add to a growing body of work demonstrating the importance of examining individual differences in tDCS response.

## Introduction

The ability to modulate brain activity *in vivo* and improve behavioral performance is crucial for advancing treatment of neurological disorders as well as combating cognitive and motor declines with age. However, providing accessible and non-invasive means to do so is a challenging objective that requires focused investigation. One promising method for non-invasive neuromodulation is transcranial direct current stimulation (tDCS). tDCS is capable of enhancing or inhibiting brain activity in cortical areas by altering neuronal firing rates and neurotransmitter concentrations (Nitsche and Paulus 2000; Stagg et al. 2009; Takano et al. 2011). Excitation of the motor cortex via tDCS consistently results in improved motor function (Rroji et al. 2015; Hashemirad et al. 2016; Saruco et al. 2017); however, the effects of tDCS on other brain areas, such as the frontal cortex, as well as the effects of tDCS on cognitive task domains, are less uniform (Jacobson et al. 2012; Mancuso et al. 2016; Imburgio and Orr 2018). Critically, this method has been shown to induce a reorganization of functional networks and impact neuroplasticity (Nitsche et al. 2008; Ruttorf et al. 2019).

### Stimulating the Midbrain with tDCS

While tDCS is capable of influencing activity across the nodes of functional networks when targeting just one cortical site, it is unclear how effective tDCS is at modulating subcortical targets. Nevertheless, recent research reveals promising evidence indicating an effect of cortical stimulation on neurotransmitter concentrations in striatal regions (Fonteneau et al. 2018; Fukai et al. 2019), as well as functional connectivity in cortico-striatal and thalamo-cortical networks (Polanía et al. 2012). We were specifically interested in the possibility of using tDCS to stimulate the dopaminergic midbrain, a subcortical system primarily composed of dopaminergic neurons in the substantia nigra and ventral tegmental area. Dopamine is important for a variety of cognitive and motor behaviors, and the dopaminergic midbrain, in particular, is implicated in reward-based decision making, learning, and motivated behavior, as well as disease pathology and aging (Howes and Kapur 2009; Bäckman et al. 2010). Modulating the midbrain using a method such as tDCS could contribute to therapeutic advances targeting behaviors and diseases involving dopaminergic dysfunction. Notably, the midbrain is both directly and indirectly linked to the prefrontal cortex via mesocortical dopaminergic pathways (Murase et al. 1993; Karreman and Moghaddam 1996; Carr and Sesack 2000; Westerink et al. 2018). Thus, prefrontal stimulation seems to cause a downstream effect in the midbrain through modulation of these dopamine-driven connections (Fonteneau et al. 2018; Fukai et al. 2019).

Combining bifrontal tDCS to the left and right dorsolateral prefrontal cortex (DLPFC) with positron emission tomography (PET), Fonteneau and colleagues (2018) recently demonstrated that stimulation induces dopamine release in the ventral striatum, which maintains connections with the midbrain through the mesolimbic dopaminergic pathway. Similarly, Fukai and colleagues (2019) demonstrated increased dopamine release in the striatum following bifrontal DLPFC stimulation, which was associated with improved accuracy on an attentional control task. This relationship between increased striatal dopamine activity and cognitive improvement after prefrontal tDCS supports the idea that cortical stimulation can be used to affect subcortical dopamine systems and associated behavioral functions.

In addition, tDCS and fMRI studies have shown that frontal stimulation increases signal intensities in the nucleus accumbens, a component of the ventral striatum (Takano et al. 2011). Clinical studies of patients with major depressive disorder have shown that prefrontal repetitive transcranial magnetic stimulation (rTMS) also increased dopamine release in the striatum, as measured by single photon emission computed tomography (SPECT) (Pogarell et al. 2006, 2007). Further, prefrontal rTMS was shown to have similar effects on dopaminergic activity in the striatum as d-amphetamine, a dopamine agonist. The developing consensus from these studies is that modulation of the meso-cortico-limbic pathway may be a possible mechanism of action behind prefrontal tDCS-induced changes in subcortical dopamine, as this pathway connects the DLPFC to the ventral striatum. In a recent resting state fMRI study, prefrontal tDCS and administration of L-Dopa, a precursor to dopamine, enhanced spontaneous neural activity in mesostriatal regions (Meyer et al. 2019). Thus, this body of research supports the idea that the meso-cortico-limbic pathway may be involved in the application of prefrontal tDCS as a moderator of subcortical dopamine activity.

### Behavioral Measures of Dopamine

While the research discussed thus far involves more direct measures of brain activity, such as PET and fMRI, more cost-effective and accessible behavioral and physiological measures have been used to indirectly estimate dopamine levels in humans. For example, behavioral paradigms using facial attractiveness ratings have been used to measure subcortical dopamine, as this particular type of task has demonstrated an association with midbrain activity (Chib et al. 2013). As demonstrated by Chib et al. (2013), stimulation of the prefrontal cortex with tDCS has been successful in increasing appraisals of facial attractiveness, presumably by way of remotely activating the dopaminergic pathways that lead to the midbrain, which then results in increased ventral midbrain activity. Thus, after administration of tDCS, facial attractiveness ratings were positively correlated with ventral midbrain activity (Chib et al. 2013). Research has also shown that facial attractiveness paradigms activate reward circuitry through frontal-striatal dopaminergic pathways (Aharon et al. 2001; Cloutier et al. 2008; Liang et al. 2010; Yu et al. 2013). In addition, reward-related brain regions have been shown to express a linear change in activity with increasing or decreasing attractiveness judgments, though some of these regions are preferentially responsive based on the subjects’ gender (Cloutier et al. 2008). Males show a stronger relationship between attractiveness judgments and activity in the orbito-frontal cortex, but other reward regions do not demonstrate sex differences. Due to the connection between facial attractiveness appraisals, reward-related brain regions, and dopamine functioning, facial attractiveness rating might serve as a proxy of subcortical dopamine levels.

Additionally, research suggests that orienting bias in visual attention is another potential proxy of subcortical dopamine functioning, as this phenomenon is related to D2 receptor asymmetries in the striatum (Tomer 2008). Orienting bias toward the left visual hemispace, specifically, is thought to arise from a right hemispheric specialization in spatial information processing. This relationship between orienting bias and dopamine is substantiated by PET evidence exhibiting that pseudoneglect, or the natural tendency to shift visual attention to the left hemispace, reflects disparities in the lateralization of dopamine systems in the striatum (Tomer et al. 2013). Additionally, differences in spatial attention have been predicted by genetic variations of the dopamine transporter gene (Newman et al. 2012; Zozulinsky et al. 2014). Thus, the degree to which an individual maintains a leftward orienting bias can be used as an additional variable to indirectly inform differences in subcortical dopamine activity.

Lastly, spontaneous eye blink rate (EBR) is also often used as a physiological proxy for dopamine levels. Baseline blink rate and tonic dopamine activity have been positively correlated in a number of studies (Karson 1983; Taylor et al. 1999; Jongkees and Colzato 2016). Further, neuroimaging work has demonstrated a link between EBR and dopamine D2 receptors (Groman et al. 2014), though more recent PET studies have failed to replicate these findings (Dang et al. 2017; Sescousse et al. 2018). However, both D1 and D2 agonists have been implicated in a dose-dependent relationship with EBR (Elsworth et al. 1991; Kleven and Koek 1996; Jutkiewicz 2004; Kaminer et al. 2011), and current research continues to employ this method as an indirect measure of tonic dopamine functioning (Byrne et al. 2019). Moreover, work by Slagter and colleagues (2010) demonstrates that EBR predicts individual differences in pseudoneglect, and, as these measures have both been related to dopaminergic activity, their relationship to each other further implicates EBR as an indirect correlate for dopamine.

### Current Study

To extend previous work, we examined the effects of prefrontal tDCS on subcortical dopamine activity using three behavioral and physiological correlates of dopamine - facial attractiveness ratings, visuospatial orienting bias, and EBR. Our primary goal was to use indirect measures of dopamine functioning to replicate previous work that used more direct measures, such as PET and fMRI. Establishing a method by which the impact of tDCS on dopamine might be measured without expensive fMRI or PET imaging would allow for a more accessible approach that future researchers can use to explore this relationship. We were also interested in evaluating the possible influence of individual variability in response to tDCS using the impact of baseline dopamine on changes in dopamine function following stimulation, through a post hoc comparison. Research suggests that the efficacy of tDCS may be related to individual differences in dopamine genes associated with various factors (e.g., attentional orienting bias, response to reward, and working memory) (Kirsch et al. 2006; Zozulinsky et al. 2014; Stephens et al. 2017); this method may, therefore, exert differing effects on behavior, depending upon an individual’s receptiveness to stimulation (Berryhill and Jones 2012; Plewnia et al. 2013; Nieratschker et al. 2015). Therefore, we used a mixed design wherein stimulation condition (active or sham) was implemented as a between-subjects variable, while behavioral and physiological data was collected both before and after stimulation as a within-subjects variable. Both before and after tDCS, we used EBR, a facial attractiveness paradigm, and a greyscales orienting bias task as a multilayered, indirect index of subcortical dopamine levels.

We predicted that active tDCS would increase midbrain dopamine levels as evidenced by a downstream impact on behavior and physiology. More specifically, we expected higher ratings on the facial attractiveness paradigm and a decrease in leftward attentional bias following active stimulation relative to sham, as well as an increase in average EBR. These hypotheses are based on the literature implicating the meso-cortico-limbic dopamine pathway in facial attractiveness appraisals, visuospatial attention, and blink rate (Aharon et al. 2001; Cloutier et al. 2008; Tomer 2008; Tomer et al. 2013; Yu et al. 2013). However, taking individual differences into account, we further predicted that baseline dopamine levels, as quantified by behavioral and physiological proxies, would impact response to tDCS. This hypothesis is based on research wherein inter-individual differences in baseline dopamine, as a product of genetic variability, were shown to influence behavioral outcomes, such as attentional bias and activation of reward systems (Kirsch et al. 2006; Zozulinsky et al. 2014; Stephens et al. 2017). Our experimental design and outline for *a priori* analyses were pre-registered on Open Science Framework before data collection began, and our *post hoc* analyses are outlined with the statistical approach for this study.

## Methods

### Participants

Thirty-four healthy young adults participated in the study. Four participants were excluded from analyses due to technical difficulties with data collection (n = 1), discomfort from stimulation (n = 1), or issues establishing an adequate connection (n = 2) as thick, curly hair can obstruct electrodes when using the particular montage employed here. As such, our final sample included thirty participants (mean age = 22.43 ± 3.15 years, 11 females). All subjects were right-handed and did not have a history of neurological illness. In addition, none of the participants were taking medication that may affect the central nervous system and none had consumed alcohol or used illicit substances on the day of the experiment. All subjects were screened according to the IRB approved exclusion criteria for tDCS studies (Nitsche et al. 2008). Participation was limited to those that had not completed other studies involving tDCS in the past. Recruitment was executed through either the Texas A&M University Psychology Subject Pool or bulk email, and all subjects were compensated $10/hour for participation. We are able to advertise paid research opportunities through the Texas A&M University Psychology Subject Pool website, and as such, participants recruited through this mechanism received compensation, as previously noted, rather than course credit for their participation. All procedures were approved by the Institutional Review Board at Texas A&M University, and written informed consent was obtained from all participants.

### Transcranial Direct Current Stimulation

We used a mixed design with stimulation condition as a between-subjects variable. Each subject participated in only one session and received either sham or active stimulation. Participants were randomly assigned to a stimulation group upon enrollment in the study. All participants were blind to the condition until the debriefing period at the end of the study. tDCS was administered using a Soterix 1×1 Low-Intensity Transcranial Electrical Stimulator and two 5cm x 7cm sponges soaked in saline (Soterix Medical, New York, NY). Six milliliters of saline solution, concentrated at 0.9% salt, was applied to each side of the sponges. An electrode was placed inside each sponge and attached to the scalp using elastic bands. The electrodes were placed on F3 and F4, the areas corresponding to the left dorsolateral prefrontal cortex (lDLPFC) and right dorsolateral prefrontal cortex (rDLPFC), respectively, according to the 10-20 measurement system. The anode was placed over the lDLPFC while the cathode was placed over the rDLPFC, in line with previous work (Fonteneau et al. 2018; Fukai et al. 2019). An initial stimulation of 1.0mA was delivered for 30 seconds in each session regardless of stimulation condition, allowing the current to break through the scalp and establish an adequate connection. Once optimal contact quality between the electrodes and the scalp was established, stimulation began and current gradually increased until the desired intensity of 2.0mA was reached. tDCS was administered at a steady 2.0mA for a period of 20 minutes during the active stimulation sessions. During sham stimulation, however, participants only experienced a current of 2.0mA at the first and last 30 seconds of the 20 minute period, while a current of 0mA was delivered for the remaining time in these sessions - this was meant to simulate the sensation of stimulation without affecting brain function. In order to assess the effectiveness of this sham procedure, questions concerning the subjects’ perception of stimulation condition were included in a final survey at the end of the session before the condition received was fully disclosed.

### Behavioral Assessments

Two behavioral tasks associated with dopaminergic activity were implemented before and after tDCS to index any changes that may occur in response to stimulation. One task was modeled after a facial attractiveness rating paradigm by Chib and colleagues (2013) where subjects were presented with a neutral-expression facial image and instructed to rate the attractiveness of that image on a scale of 1 to 7 (Figure 1), where “1” indicated a low attractiveness score while “7” indicated a high attractiveness score. These ratings were made using a computer keyboard. Participants completed 72 trials both before and after tDCS for a total of 144 trials. The faces were displayed on the computer screen until the participant made a response, at which point the next stimulus would be presented after an inter-stimulus interval of 1000 ms. The images were taken from the Chicago Face Database, Version 2.0.3 (Ma et al. 2015), which includes normative ratings of attractiveness for each face. The faces were grouped and randomized such that each set, both before and after stimulation, was equal in mean attractiveness. The image sets were also generated to include an equal percentage of male and female faces and an equal proportion of Asian, Black, Latinx, and Caucasian faces. We measured the difference in average attractiveness ratings between pre-tDCS and post-tDCS performance.

The greyscales paradigm (Figure 2) was used to assess the degree of leftward visuospatial bias exhibited by each individual. The task parameters were consistent with those used by Tomer and colleagues (2013). Two rectangles containing a black to white gradient were presented on a computer screen for 5000 ms and each subject was instructed to choose the rectangle that appeared darker overall. After 5000 ms passed, the rectangles were whited-out. The next gradient stimulus was not presented until a response was made. If the participant made a response before the gradients became completely white, the trial was ended and the next stimulus appeared after an inter-stimulus interval of 1500 ms. Subjects were instructed to press “B” to indicate that they chose the bottom rectangle or “T” to indicate that they chose the top rectangle. Following an initial practice portion of 12 trials, the task included 144 experimental trials (Tomer et al. 2013) and was administered both before and after stimulation. In the first half of the trials, a difference in luminosity was present, but in the second half there was no difference in darkness between the rectangles. Leftward attentional bias was measured as the proportion of trials where the subject chose the rectangle with a gradient beginning on the left side as the darker stimulus overall when both rectangles were in fact the same. In addition, looking only at error trials where subjects failed to accurately respond, attentional bias was calculated for trials in which a difference in darkness was indeed present between rectangles. However, the latter calculation was a supplementary assessment of orienting bias. An increase in dopamine following active tDCS would result in a reduction of leftward attentional bias after stimulation, relative to before. Therefore, dopaminergic modulation was indirectly indexed by the difference in leftward attentional bias between pre- and post-tDCS performance.

**Figure 2.**
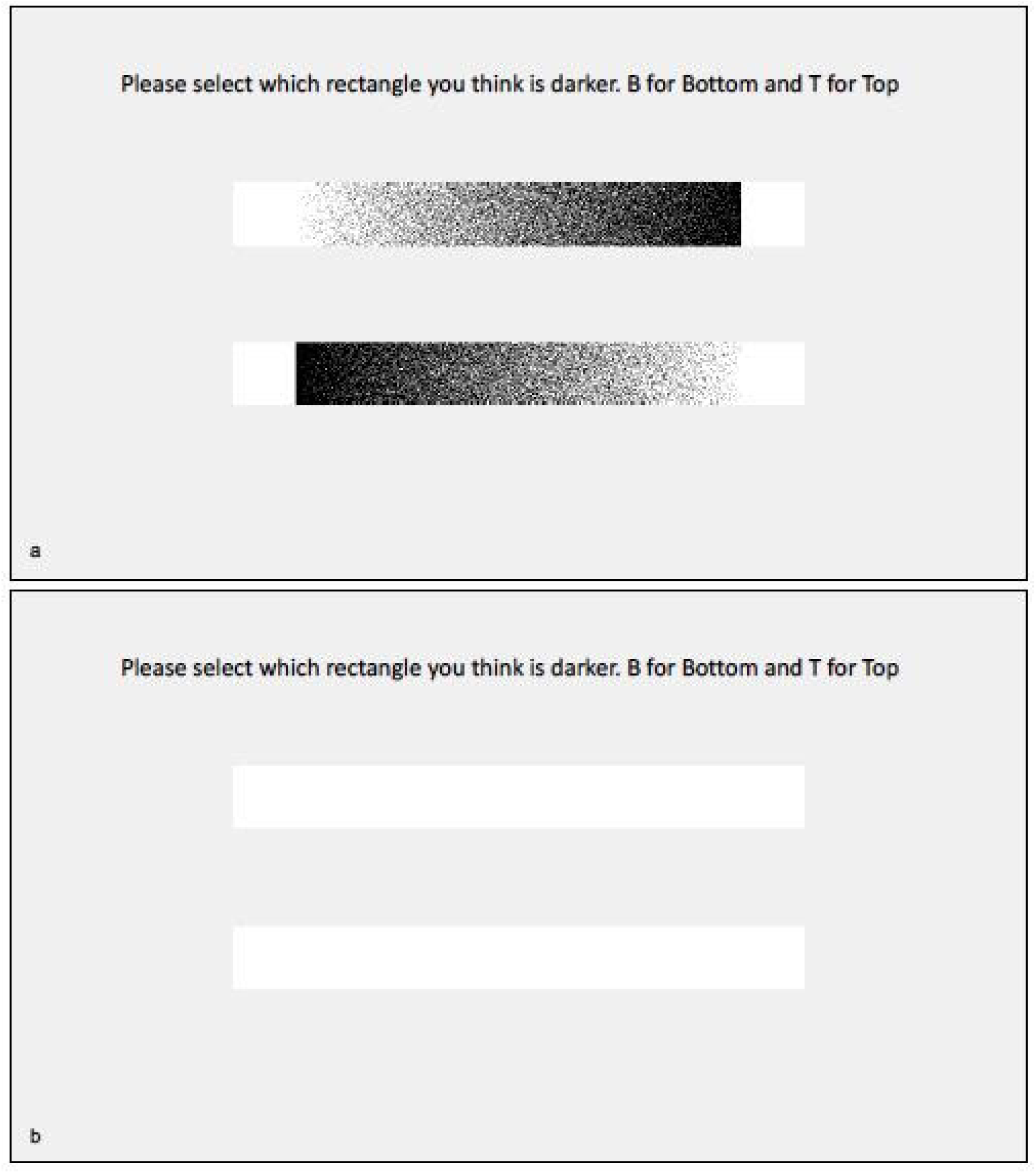
Greyscales paradigm. Participants were instructed to choose the rectangle that appeared darker overall using the “B” key for bottom and the “T” key for top on the computer keyboard (a). After 5000 ms passed, the rectangles were whited-out until the participant made a response (b). The task consisted of 144 trials, administered both before and after tDCS.

### Physiological Assessment

In addition to the behavioral assessments, spontaneous eye blink rate (EBR) was recorded as a physiological marker of dopaminergic activity both before and after the stimulation period. Using Ag/AgCl electrodes and a BIOPAC EOG100C Electrooculogram Amplifier, eye movements were recorded upon securing two receiving electrodes around the right eye and one ground electrode between the eyebrows (BIOPAC Systems, Inc., Goleta, CA). Participants were instructed to focus on, but not stare at, a fixation cross without engaging in any activity while their natural blink rate was recorded for a 5-minute period. This was preceded by a 30-second baseline recording where markings distinguishing between eye movements and eye blinks were made to assist in subsequent data analysis. This procedure was repeated again after tDCS, and blinks were manually counted by two separate raters before a blink rate average per 30 second interval was calculated for each subject. Inter-rater reliability for the total number of blinks during each 5 minute recording was very high (*r* = 0.99). We measured the difference in average EBR after tDCS, relative to before, to gauge any changes in tonic dopamine activity as a result of stimulation.

### Procedure

Upon arrival, participants were given a brief overview of the study procedures and completed a consent form. Participants then performed the greyscales and facial attractiveness tasks to provide baseline behavioral assessments. The order of these tasks was counterbalanced across all subjects. Once both tasks were completed, EBR setup began and recordings were conducted for a period of 5 minutes while the participant fixated on a cross as instructed. Following this period, the EBR electrodes were disconnected from the amplifier but left attached to the subject during tDCS to keep placements consistent and minimize variability between pre-tDCS and post-tDCS recordings. Participants then received 20 minutes of either sham or active stimulation, based on random stimulation group assignment. After tDCS was completed, the equipment was removed and a brief demographic survey was administered. EBR electrodes were then reconnected to the amplifier to complete another 5 minute period of eye blink recordings before a second round of the same behavioral tasks took place. In addition, a questionnaire regarding sensations from tDCS was administered to collect information on the subjects’ experience. Finally, each participant was debriefed on the purpose of the study and the stimulation condition was revealed.

### Data Analyses

All analyses not listed as exploratory were pre-registered, along with the study design, on Open Science Framework (https://osf.io/gmnpc). Independent samples t-tests were used to assess the effects of stimulation condition on each dopamine correlate individually, where the dependent variable was change in the measure (post stimulation - pre stimulation). Independent samples t-tests were also used to compare measures across stimulation groups prior to stimulation to ensure that any difference in the post-stimulation measures was not due to group differences prior to stimulation. Finally, Bayes factors (BF_01_) were calculated for each test to quantify the likelihood of a null result (no effect of tDCS) vs. the pre-registered a-priori hypotheses. These Bayes factors represent how likely the null hypothesis is in comparison to the alternative. Following previous work (Biel and Friedrich 2018), a Bayes factor of greater than or equal to 3, meaning the null is 3 times more likely than the alternative, was considered to conclusively support a null result. Because mean facial attractiveness ratings were skewed towards the lower end of the rating scale (*M* = 3.18, *Max* = 4.72, *Min* = 1.08), attractiveness ratings were max-normalized following Chib and colleagues (2013). All subsequent references to attractiveness ratings refer to these max-normalized ratings. Additional analyses regarding the gender of the faces and participants’ sexual preference can be found in the Supplementary Materials.

Previous work relating leftward bias on the greyscales task to dopamine systems has focused primarily on bias in trials on which there is no difference in darkness between stimuli (Tomer 2008; Slagter et al. 2010; Tomer et al. 2013; Zozulinsky et al. 2014). However, previous work has also shown that on greyscales trials in which there is a difference between the darkness of stimuli, the leftward bias on error trials is correlated with leftward bias on trials with equally dark stimuli and might also be related to striatal dopamine (Tomer 2008; Tomer et al. 2013). Therefore, we examined the effect of stimulation on bias in trials where there is no difference between stimuli, but bias in trials on which there is a difference is examined in the supplementary materials. Leftward bias was calculated as the percent of trials on which the rectangle that was darker on the left was chosen. On trials in which there was no difference in darkness between the two stimuli, participants displayed a leftward bias prior to the stimulation period (*M* = 64.58%, *SD* = 20.38%), after the stimulation period (*M* = 63.89%, *SD* = 25.41%), and across both time points (*M* = 64.24%, *SD* = 22.84%).

Sham stimulation effectiveness was first assessed by examining the survey question that asked which condition participants believed they were in. Results are reported descriptively in Table 1 and are analyzed formally in the supplemental materials.

**Table 1.**
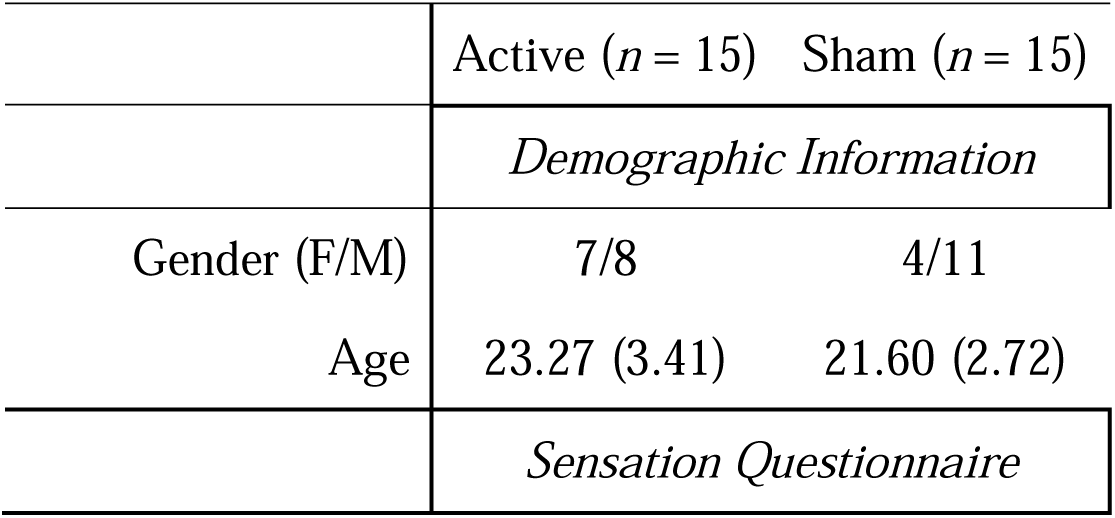
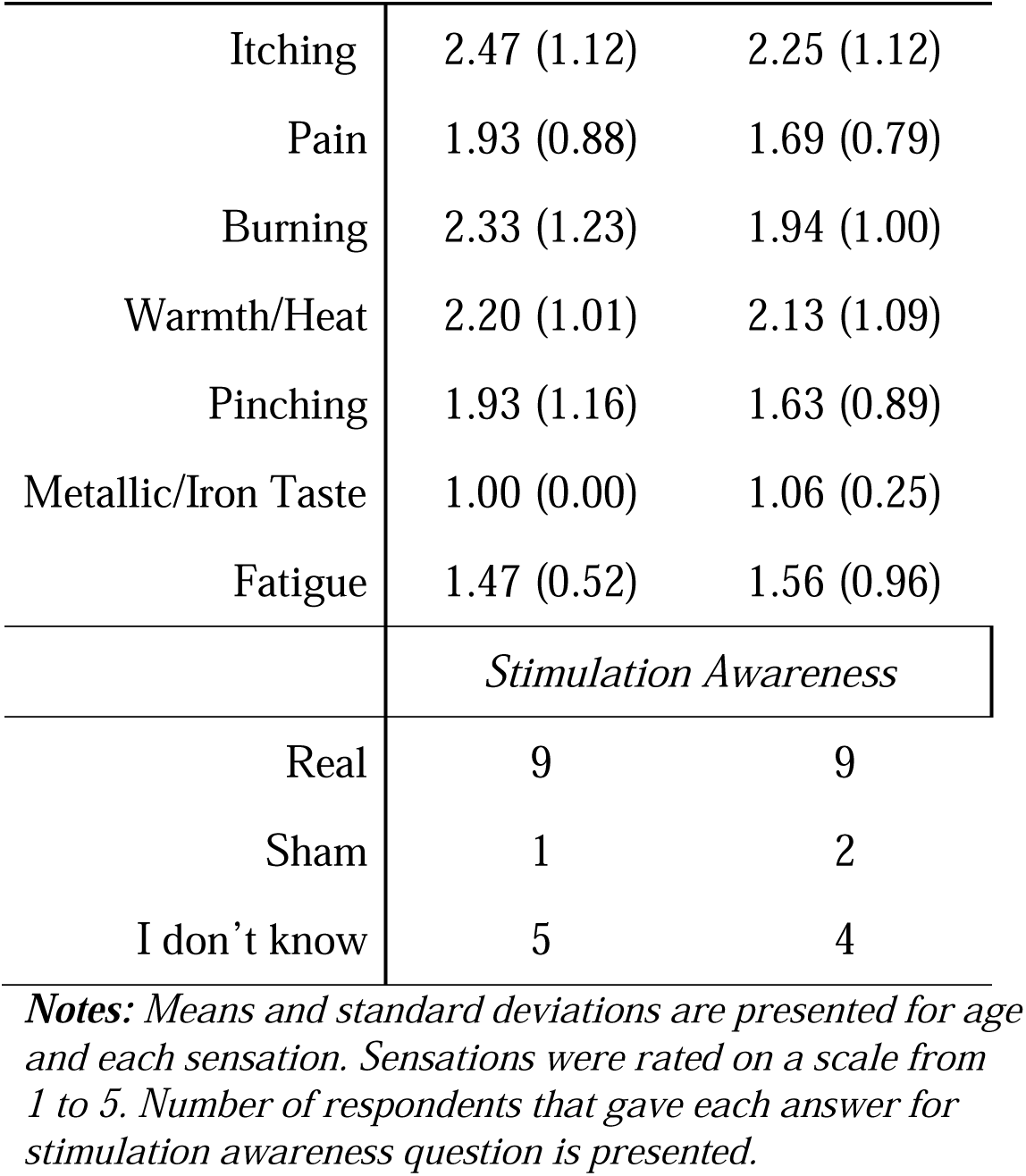
Demographics and Placebo Effectiveness

| | Active ($n = 15$) | Sham ($n = 15$) |
| --- | --- | --- |
|  | *Demographic Information* |  |
| Gender (F/M) | 7/8 | 4/11 |
| Age | 23.27 (3.41) | 21.60 (2.72) |
|  | *Sensation Questionnaire* |  |
| Itching | 2.47 (1.12) | 2.25 (1.12) |
| Pain | 1.93 (0.88) | 1.69 (0.79) |
| Burning | 2.33 (1.23) | 1.94 (1.00) |
| Warmth/Heat | 2.20 (1.01) | 2.13 (1.09) |
| Pinching | 1.93 (1.16) | 1.63 (0.89) |
| Metallic/Iron Taste | 1.00 (0.00) | 1.06 (0.25) |
| Fatigue | 1.47 (0.52) | 1.56 (0.96) |
|  | *Stimulation Awareness* |  |
| Real | 9 | 9 |
| Sham | 1 | 2 |
| I don't know | 5 | 4 |

*Notes: Means and standard deviations are presented for age and each sensation. Sensations were rated on a scale from 1 to 5. Number of respondents that gave each answer for stimulation awareness question is presented.*

As exploratory analyses, we were interested in whether the effect of tDCS on one behavioral measure might depend upon an individual’s baseline (pre-stimulation) dopamine as measured by the three tasks. We aimed to aggregate scores across the three tasks to operationalize baseline dopamine. However, to test whether the three pre-stimulation measures (attractiveness ratings, leftward bias, and EBR) were related enough to aggregate across, we ran a factor analysis to confirm that the three measures loaded onto a single factor. Then, three regressions were conducted in which the DV was change in one measure (post-stimulation minus pre-stimulation), and the IVs were stimulation condition and a composite formed using the other two measures (formed by converting the measures to z-scores and averaging across them), hereafter referred to as a baseline dopamine composite. For leftward bias scores, z-scores were sign-flipped prior to averaging, as a lower leftward bias is indicative of greater dopamine. We purposefully did not, for example, include baseline EBR in the composite if change in EBR was the DV to avoid issues that might arise from including the same measure in both. Thus, the composite for each regression was only composed of two of the three measures. However, conclusions did not change when all three measures were included in the baseline dopamine composite score.

All p-values reported in the exploratory analyses section are false-discovery rate (FDR) corrected (*p*_*ADJ*_). Resulting p-values that remained below .05 following FDR correction were considered significant. Additional analyses that led to the conclusion reached in the ‘Exploratory Analyses’ section may be found in the Supplementary Materials and were included in the multiple comparisons corrections.

All analyses were conducted using R 3.6.2 (R Core Team 2019) and JASP 0.9.2 (JASP Team 2019).

## Results

### Pre-registered Analyses

Prior to stimulation, there was no significant difference across stimulation groups in leftward bias on the greyscales task *t*(27.90) = 0.50, *p* = .62. There was no effect of stimulation condition on change in leftward bias, *t*(19.72) = 1.27, *p* = .22, *d* = 0.47 (see Fig. 3A). Bayesian analyses did not yield strong conclusions favoring the null or the alternative hypotheses (*BF*_*01*_ = 0.92). However, the active stimulation group exhibited a greater decrease in bias following stimulation (*M* = -5.19%, *SD* = 24.78%) than the sham stimulation group (*M* = 3.80%, *SD* = 11.45%), in line with predictions.

**Figure 3.**
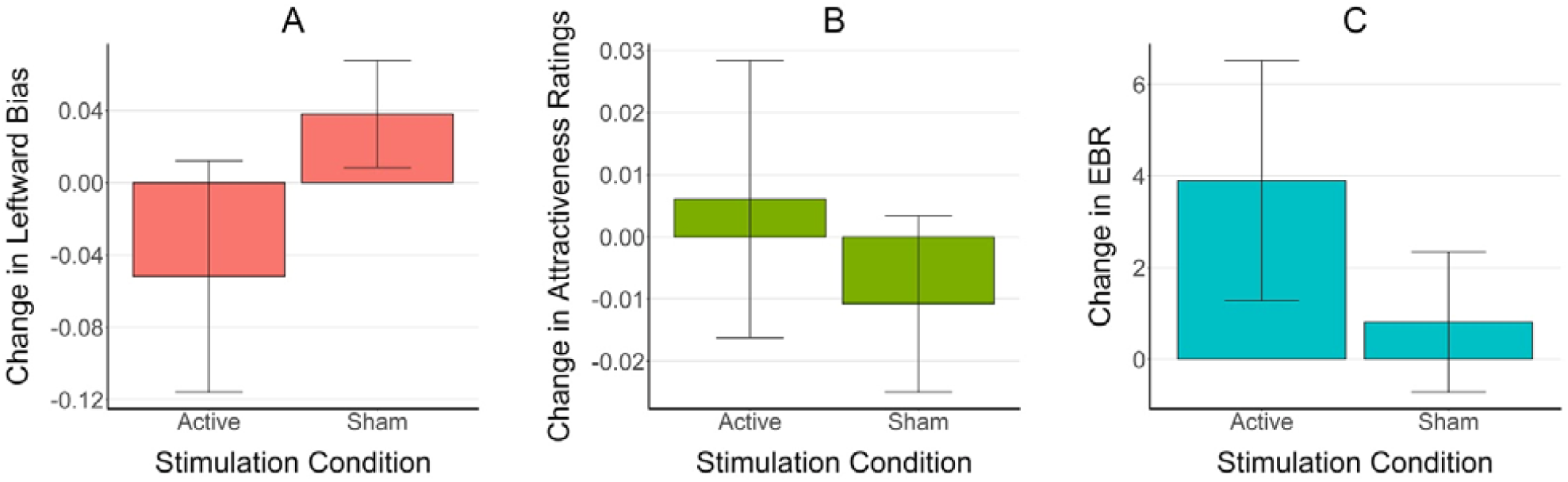
Pre-registered analyses assessing the effects of tDCS on changes in leftward bias on the greyscales task (A), attractiveness ratings (B), and EBR (C). All changes were calculated as poststimulation minus pre-stimulation. No effects of stimulation were significant, although all effects were in the predicted direction. Error bars represent standard error.

Prior to stimulation, there was no significant difference between stimulation groups in attractiveness ratings in response to faces *t*(24.45) = 0.05, *p* = .96. There was no effect of stimulation condition on change in attractiveness ratings, *t*(23.73) = 0.64, *p* = .53, *d* = 0.23 (see Fig. 3B). The null hypothesis was more likely than the alternative, but not conclusively so (*BF*_*01*_ = 1.76).

Prior to stimulation, there was no significant difference between stimulation groups in EBR, *t*(29.97) = 0.31, *p* = .76. There was no effect of stimulation on change in EBR (*t*(24.31) = 1.19, *p* = .25, *d* = 0.42; see Fig. 3C). Bayesian analyses were inconclusive - they did not strongly support the null or the alternative hypothesis (*BF*_*01*_ = 1.22). The direction of the effect was, however, in line with predictions; there was a larger increase in EBR following active stimulation (*M* = 3.90, *SD* = 10.15) compared to sham stimulation (*M* = 0.81, *SD =* 5.92).

### Exploratory Analyses

Although stimulation did not significantly affect any of the measures across the entire sample, effects were in the expected direction. We reasoned that this might, in part, be due to individual variability in response to tDCS. Because some prior work has found variability in tDCS response to depend upon dopamine-related genes (Plewnia et al. 2013; Nieratschker et al. 2015; Shivakumar et al. 2015; Stephens et al. 2017), we examined an individual’s baseline dopamine (operationalized as an aggregate of our measures) prior to stimulation as a possible predictor of tDCS effects.

Prior to aggregating across the three measures, we conducted an exploratory factor analysis to assess whether the measures all loaded onto a single factor. An examination of the scree plot (Fig S1) indicated that the single factor solution was the only model that yielded eigenvalues above one. The one factor solution, which accounted for 41% of the variance, was indeed the best-fitting solution (factor loadings are listed in Table S1).

When pre-stimulation greyscales bias and attractiveness ratings were aggregated to assess overall baseline dopamine, there was a significant interaction between stimulation condition and composite baseline dopamine in predicting change in EBR (β = -1.13, *p*_*ADJ*_ = .02, *R*^*2*^_*adj*_ = .29; see Fig. 4). Within participants that received active stimulation, change in EBR was positively correlated with the composite baseline dopamine score (β = .58, *p*_*ADJ*_ = .04). Within participants that received sham stimulation, change in EBR was negatively correlated with composite baseline dopamine score (β = -.61, *p*_*ADJ*_ = .04). Together, these results suggest that tDCS was more effective in increasing EBR for participants with higher levels of dopamine prior to stimulation. There was no significant interaction between stimulation condition and composite baseline dopamine score in predicting change in facial attractiveness ratings (β = .47, *p*_*ADJ*_ = .30, *R*^*2*^_*adj*_ = -.02) or leftward bias (β = -.42, *p*_*ADJ*_ = .34, *R*^*2*^_*adj*_ = .01).

**Figure 4.**
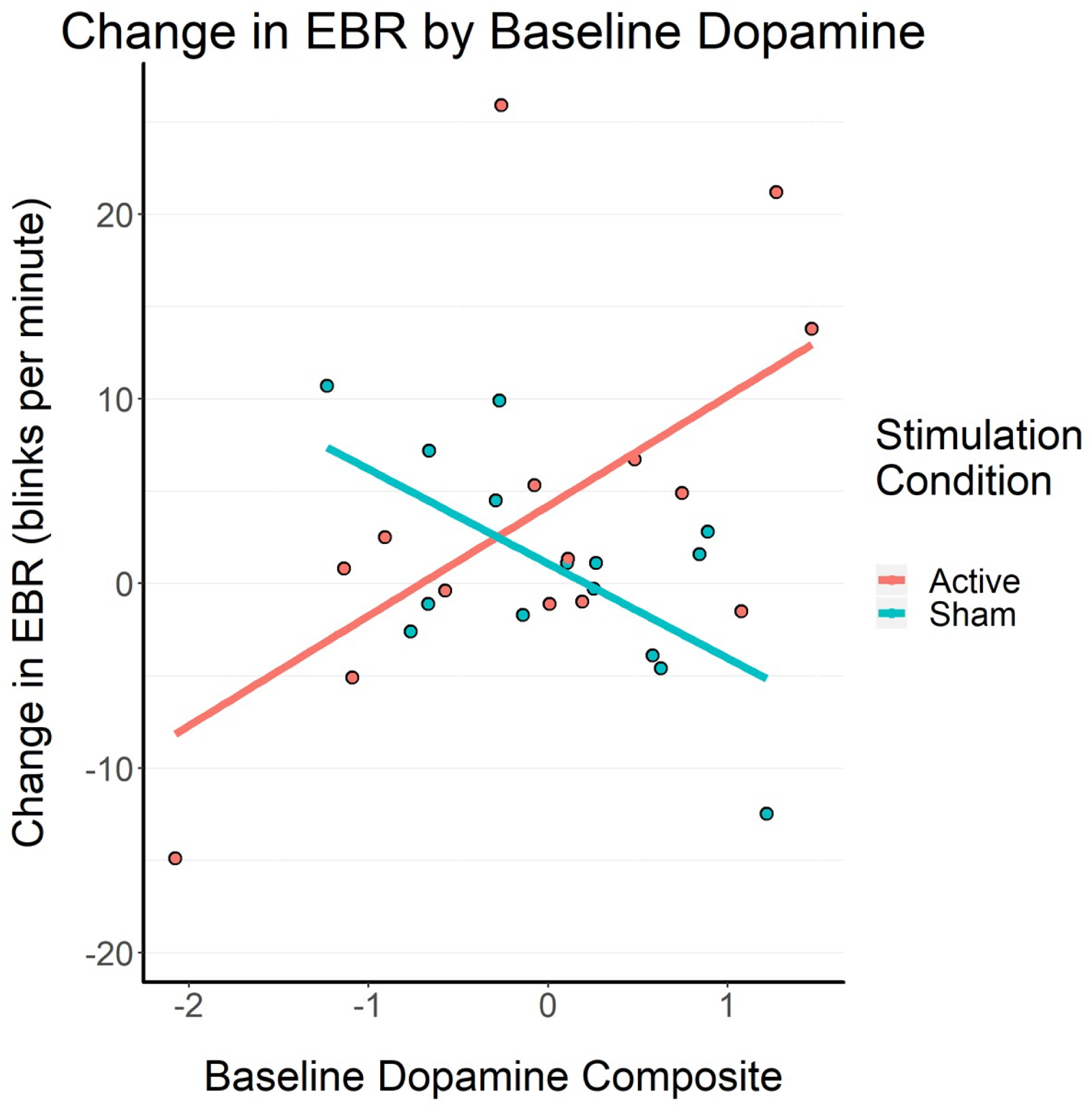
Interaction between baseline dopamine composite score (formed using pre-stimulation leftward bias and attractiveness ratings) and change in EBR following stimulation. Change in EBR following the stimulation period was positively correlated with baseline dopamine for the active stimulation participants, but negatively correlated for the sham stimulation group.

## Discussion

The current work sought to replicate previous PET studies that report an increase in striatal dopamine following tDCS using an anode over F3 and a cathode over F4 by using the same montage and three indirect measures of dopamine. We failed to find support for our preregistered hypotheses - that tDCS would increase EBR, increase facial attractiveness ratings, and decrease leftward visuospatial bias, all theoretically indicative of an increase in dopamine. In exploratory analyses, we found that the effect of tDCS on EBR was significantly moderated by an individual’s pre-stimulation dopamine, as measured by their scores on the facial attractiveness and greyscales tasks. Participants with scores indicating higher baseline dopamine exhibited a greater increase in EBR as a result of active stimulation and those with lower baseline dopamine scores experienced no change or a decrease in EBR. In the sham stimulation group, this relationship was reversed.

Two recent studies using the same tDCS montage as the current work reported an increase in striatal dopamine as measured by PET (Fonteneau et al. 2018; Fukai et al. 2019). While we were not able to detect the same change in our sample using indirect measures, all effects were in the predicted direction. Further, the effect sizes of the changes were moderate, particularly for changes in EBR (*d* = .42) and leftward bias (*d* = .47), and Bayesian analyses indicated that the null hypothesis (that tDCS did not affect the measures) was not conclusively more likely than the alternative (*BF*_*01*_s ≤ 1.76). Therefore, it is possible that the indirect measures are not sensitive enough to detect any tDCS-induced changes in dopamine in a between-subjects sample of this size (*N* = 30). In line with this idea, Fukai and colleagues (2019) estimated an effect size of 1.27 using PET measures of dopamine, a substantially larger effect than those reported here. While our pre-registered sample size can be considered standard in the tDCS literature - one meta-analysis of tDCS studies reported an average sample size of 21 (Nilsson et al. 2015) and a number of tDCS studies published in the last year have utilized samples within the same range as ours (Berryhill and Jones 2012; Wolkenstein and Plewnia 2013; Gbadeyan et al. 2016; Savic et al. 2017; Fonteneau et al. 2018) - a retrospective power analysis in G*Power 3 (Faul et al. 2007) using the effect sizes reported here and a power level of 80% indicates that a sample of 172 participants would be necessary to detect any effects in a between-subjects design. Because this is such a large sample size for a typical tDCS study, collecting an ideal sample size would be best suited for a concerted group effort.

It is possible that indirect measures of dopamine, such as EBR, are insufficiently sensitive to reliably detect changes in dopamine within small, healthy samples. However, a recent review reported that the majority of studies examining the effect of dopamine agonists on EBR in humans reported no effect, and that all of the studies used samples of less than 40 participants (Jongkees and Colzato 2016). Meanwhile, much of the widely cited literature linking EBR (the most widely used of the three indirect measures in this study) to dopamine in humans is related to clinical populations - individuals with Parkinson’s disease exhibit decreased EBR (Fitzpatrick et al. 2012), individuals with schizophrenia exhibit increased EBR (Chen et al. 1996), and chronic cocaine users exhibit decreased EBR (Colzato et al. 2008). This pattern in the literature may indicate that, in smaller samples, EBR is only sensitive enough to detect dopamine differences in clinical populations rather than intervention-induced changes in healthy populations. No work to date has examined changes in facial attractiveness ratings after administration of a dopamine agonist, and the few studies finding changes in visuospatial bias in response to drug manipulations rely on case studies of clinical patients (Fleet et al. 1987; Geminiani et al. 1998), rather than larger healthy samples.

Of course, it is also possible that the relationships between indirect dopamine measures and direct dopamine measures are still nonsignificant in larger healthy samples. However, as demonstrated in the current work, sample sizes considered standard in tDCS and PET literature are still likely insufficient to demonstrate (using Bayesian analyses) that no relationship between direct and indirect measures is more likely than a weak relationship (Minarik et al. 2016; Medina and Cason 2017).Therefore, even if the measures used in the current work are unrelated to dopamine in healthy human samples, the field would greatly benefit from an examination of this relationship in larger sample sizes.

It should be noted that Tomer and colleagues (2013) found a relationship between dopamine asymmetry in the striatum and visuospatial bias, rather than the absolute value of dopamine in either hemisphere. Further, EBR is moderately correlated with visuospatial bias (Slagter et al. 2010), which might suggest that EBR is more sensitive to dopamine asymmetry than the absolute value of dopamine in the striatum. Future PET work should examine this relationship in addition to the relationship between EBR and the magnitude of dopamine in the striatum in general.

The current work failed to replicate a previous study that found an increase in facial attractiveness ratings in a sample of 40 participants following prefrontal tDCS stimulation (Chib et al. 2013). This might be due to differences in stimulation montage; while the current work employed a bilateral DLPFC montage, Chib and colleagues placed their anode over the ventromedial prefrontal cortex. Imaging-based parcellations of the striatum indicate that the medial PFC and the DLPFC project to different areas of the striatum (Choi et al. 2012; Pauli et al. 2016). Meanwhile, fMRI studies have consistently linked facial attractiveness ratings to activation in the striatum and medial areas of the prefrontal cortex (Aharon et al. 2001; Smith et al. 2010; Yu et al. 2013), targeted by the montage used by Chib and colleagues. However, of the three measures used in the current work, the relationship between facial attractiveness ratings and dopamine is the least supported; no work to date has examined the relationship between this indirect measure and dopamine using PET.

Exploratory analyses indicated that stimulation may be more effective in increasing EBR for participants with higher baseline dopamine levels than participants with lower baseline dopamine levels. The positive correlation between baseline dopamine measures and change in EBR is in line with a previous study which found greater effects of a dopamine agonist on EBR within primates that had more dopamine receptors prior to drug administration (Groman et al. 2014), as well as recent human work which found a greater effect of L-dopa on cognitive control for participants with greater pre-treatment dopamine levels (Lee et al. 2019). The relationship between dopamine and stimulation effect might, in part, explain some reports that tDCS affects healthy samples and clinical samples differently (Dedoncker et al. 2016). However, even in healthy samples, dopamine receptor density (Farde et al. 1995) and the rate of dopamine synthesis (Laakso et al. 2005) varies across individuals substantially. Therefore, the relationship might contribute to the variability in tDCS effects in healthy samples, further necessitating the use of larger samples to detect group effects.

Notably, however, the exploratory results in the current work stand in contrast to PET work examining the effect of the same tDCS montage, which concluded that there was no relationship between pre-stimulation dopamine and the effects of tDCS (Fonteneau et al. 2018). There are a number of possible reasons for the discrepancy between the findings reported by Fonteneau and colleagues and those reported in our exploratory analyses. Due to the binding properties of the radiotracer used, the PET study focused primarily on baseline dopamine at D2 receptors in the striatum (as is standard in the field). The indirect measures used in the current work might additionally index dopamine at other receptor subtypes - previous work indicates that EBR might be related to D1 receptor activity as well as D2 receptor activity, for example (Elsworth et al. 1991; Kleven and Koek 1996; Jutkiewicz 2004). Indeed, recent work by Meyer and colleagues (2019) compared fMRI activation patterns from genes related to dopamine receptor subtypes to fMRI activation patterns following prefrontal tDCS. Although the montage differed from the montage in the current work, the results suggested that tDCS might affect dopamine at D3 receptors in addition to D2 receptors; future work should similarly examine how baseline dopamine at subtypes other than D2 might modulate tDCS.

The indirect measures in the current work might capture dopamine in areas outside of the striatum, such as the prefrontal cortex, also not examined in PET work. The idea that baseline prefrontal dopamine might affect an individual’s response to tDCS is not new; a number of previous studies have concluded that COMT, a gene that regulates dopamine metabolism in the prefrontal cortex, can modulate the effects of tDCS (Plewnia et al. 2013; Nieratschker et al. 2015; Shivakumar et al. 2015; Stephens et al. 2017). However, the idea that the measures used in the current work might be related to COMT or prefrontal dopamine is highly speculative, as no previous work has examined relationships between the measures used here and COMT.

Given that direct measures of dopamine found no effect of baseline dopamine on the effect of tDCS on the striatum, and that the indirect measures used in the current work are comparatively weakly related to dopamine at best, the conclusions reported here regarding the effect of baseline dopamine should be interpreted with caution. However, the variability in individual response to tDCS seen in our sample is consistent throughout tDCS literature (Wiethoff et al. 2014; Li et al. 2015; Chew et al. 2015; Laakso et al. 2019), and the current results add to an existing body of work that emphasizes the importance of understanding these individual differences.

### Limitations and Future Directions

The main limitation of the current work is the lack of direct measures of dopaminergic activity. While the measures used in the current paper have been consistently linked to dopamine systems and reward-related behavior (Karson 1983; Elsworth et al. 1991; Kleven and Koek 1996; Taylor et al. 1999; Aharon et al. 2001; Fink et al. 2002; Jutkiewicz 2004; Tomer 2008; Smith et al. 2010; Kaminer et al. 2011; Tomer et al. 2013; Yu et al. 2013), recent PET studies in humans have called the relationship between EBR (the most commonly used of those presented here) and dopamine into question (Dang et al. 2017). Further, even if these measures do reliably index striatal dopamine, they do not provide clear information about the mechanism by which prefrontal tDCS modulates striatal dopamine. Future studies should build upon the work of Fonteneau and colleagues (2018) and Fukai and colleagues (2019) by examining the effects of tDCS stimulation on D1-like receptors in the striatum and dopamine receptors in the prefrontal cortex using direct measures as well as in-depth analyses of its effects on striatocortical network function rather than solely striatal function.

The conclusions drawn in the current work are also limited by the small sample size. The sample size was determined and pre-registered using other tDCS studies as a reference point. Retrospective power analyses, however, indicate that much larger samples would be needed to detect an effect of tDCS on these indirect measures. Further, we were not able to provide support for a lack of effect of tDCS on any of our measures. Future work examining the effects of tDCS on these measures, as well as the relationship between these measures and dopamine, should aim to utilize larger samples to yield more conclusive results.

Our sample size is particularly small for individual difference analyses (although these were not planned during preregistration of the study). Because tDCS often yields very variable results, and a number of things can influence an individual’s response to tDCS including skull thickness, amount of CSF, and genetics (Nieratschker et al. 2015; Opitz et al. 2015), future work should expand on individual differences in stimulation response using larger sample sizes than those that are considered the standard in tDCS literature currently.

Understanding how tDCS affects dopamine, and how baseline dopamine levels affect an individual’s response to stimulation in general, can aid researchers that hope to modulate behavior as well as clinicians that hope to treat disorders using tDCS. Previous work examining the effectiveness in treating depression using tDCS, for example, has yielded conflicting results (Shiozawa et al. 2014); baseline dopamine differences across samples might account for some of these differences. Moreover, a previous meta-analysis concluded that tDCS was effective in modulating executive function in clinical, but not healthy, populations (Dedoncker et al. 2016); dopamine differences between the two groups may account for this difference as well. In any case, controlling for dopamine levels at baseline might help identify individuals for which tDCS is maximally effective, or not effective, in modulating behavior and treating disorders.

## Declarations

### Data and Code Availability

All data, analysis scripts and task code are publicly available on OpenScience Framework (https://osf.io/zhjys/).

### Author Contributions

Michael J. Imburgio collected and analyzed data, wrote the second half of the manuscript and contributed to study design. Hannah K. Ballard collected data and wrote the first half of the manuscript. Astin C. Cornwall oversaw EBR data collection and coded behavioral tasks. Darrell

A. Worthy contributed to manuscript preparation. Jessica A. Bernard and Joseph M. Orr contributed to manuscript preparation and study design and conceived the study.

### Informed Consent

All research was performed in accordance with IRB regulations and informed consent was obtained from all participants.

