## Supplemental Materials for "Effects of prefrontal tDCS on dopamine-mediated behavior and psychophysiology"

*Placebo effectiveness*

A chi square test of independence was used to examine whether a participants’ stimulation condition affected their answer. Next, participants’ ratings of each sensation during stimulation were analyzed as a function of stimulation condition using independent t-tests.

Participants from both stimulation conditions were equally likely to respond ‘Real’ (*n*_sham_ = 10, *n*_active_ = 9), ‘Placebo’ (*n*_sham_ = 2, *n*_active_ = 1), and ‘I don’t know’ (*n*_sham_ = 4, *n*_active_ = 5) when asked which condition they were in, 𝜒^2^(2) = 0.47, *p* = .79. There was also no effect of stimulation condition on participants’ ratings of any sensations during stimulation (all *p*s > .33), indicating that the sham stimulation procedure was effective and participants were blind to their stimulation condition.

*Pre-registered analyses of leftward bias*

On trials in which there was a difference in darkness between the two stimuli, participants chose the darker stimulus with 78.54% accuracy (*SD* = 17.12%). Leftward bias for incorrect trials was only calculated for participants that responded with less than 90% accuracy (*N* = 20), in line with previous work {{CITATION}}. Participants displayed a leftward bias on incorrect trials prior to the stimulation period (*M* = 64.51%, *SD* = 19.87%), after the stimulation period (*M* = 59.51%, *SD* = 21.97%), and overall (*M* = 62.01%, *SD* = 20.83%). There was no significant difference across stimulation groups in leftward bias on error trials prior to stimulation, *t*(17.90) = 0.34, *p* = .74. The effect of stimulation was assessed using an independent samples t-test (DV: change in error trial leftward bias, post-stimulation minus pre-stimulation; IV: stimulation condition). There was no significant effect of stimulation condition on change in error trial leftward bias, *t*(16.11) = 0.65, *p* = .52, *d* = .30.

*Pre-registered analyses of facial attractiveness ratings*

Participant’s self-reported sexual preferences were re-coded such that higher numbers always meant greater sexual attraction to females. To examine whether stimulation only affected ratings of faces of participants’ preferred gender, a regression model was fitted using change in attractiveness ratings as the dependent variable and face gender, participant sexual preference, and stimulation condition as predictors. There were no significant predictors in the model (all *p*s > .20), suggesting that even controlling for whether a participant was rating a face that matched their preferred gender, stimulation did not affect attractiveness ratings.

*Exploratory analyses*

As initial exploratory analyses examining the interaction between a participants’ baseline task performance (indicators of baseline dopamine) and the effect of stimulation, six regression models were fitted. For each model, change in one behavioral measure was the dependent variable. The independent variables were baseline (pre-stimulation) performance on a separate task and stimulation condition. Because these analyses were not planned, an FDR-correction was applied to the resulting p-values, taking into account the three exploratory regressions reported in the manuscript as well as the six exploratory regressions below.

There was no significant interaction between baseline EBR and stimulation condition (*p_ADJ_* = .87) nor between baseline attractiveness ratings and stimulation condition (*p_ADJ_* = .21) in predicting change in greyscales bias. Additionally, there was no significant interaction between baseline EBR and stimulation condition (*p_ADJ_* = .69) nor between baseline greyscales bias and stimulation condition (*p_ADJ_* = .21) in predicting change in facial attractiveness ratings.

However, stimulation condition and baseline attractiveness ratings significantly interacted to predict change in EBR (β = -0.94, *p_ADJ_* = .03). Within the active stimulation condition, participants who made higher attractiveness ratings prior to stimulation displayed a greater increase in EBR following stimulation (*r* = .70, *p_ADJ_* = .02), whereas this relationship was weaker and in the opposite direction in the sham stimulation group (*r* = -.25, *p_ADJ_* = .38). The direction of these relationships indicated that baseline dopamine was positively related to change in EBR within the active stimulation group. Additionally, stimulation condition and baseline greyscales bias interacted to predict change in EBR, although following multiple comparisons corrections the significance reached only a trend level (β = 0.84, *p_ADJ_* = .06). However, the direction of these relationships also indicated that greater baseline dopamine predicted a greater change in EBR in the active stimulation group while the opposite was true in the sham stimulation group; greater changes in EBR were associated with a nonsignificant decreased leftward bias in the active stimulation group (*r* = -.31, *p_ADJ_* = .87), while this relationship was reversed in the sham stimulation group (*r* = .67, *p_ADJ_* = .21).

*Table S1. Factor Loadings by Measure*

|  | Factor Loading |
| --- | --- |
| Facial Attractiveness Rating | 1.00 |
| Leftward Visuospatial Bias | -0.42 |
| Eyeblink Rate | 0.26 |

***Notes:*** *All measures are pre-stimulation scores. A negative leftward bias loading was expected as decreased bias is hypothesized to indicate greater dopamine.*

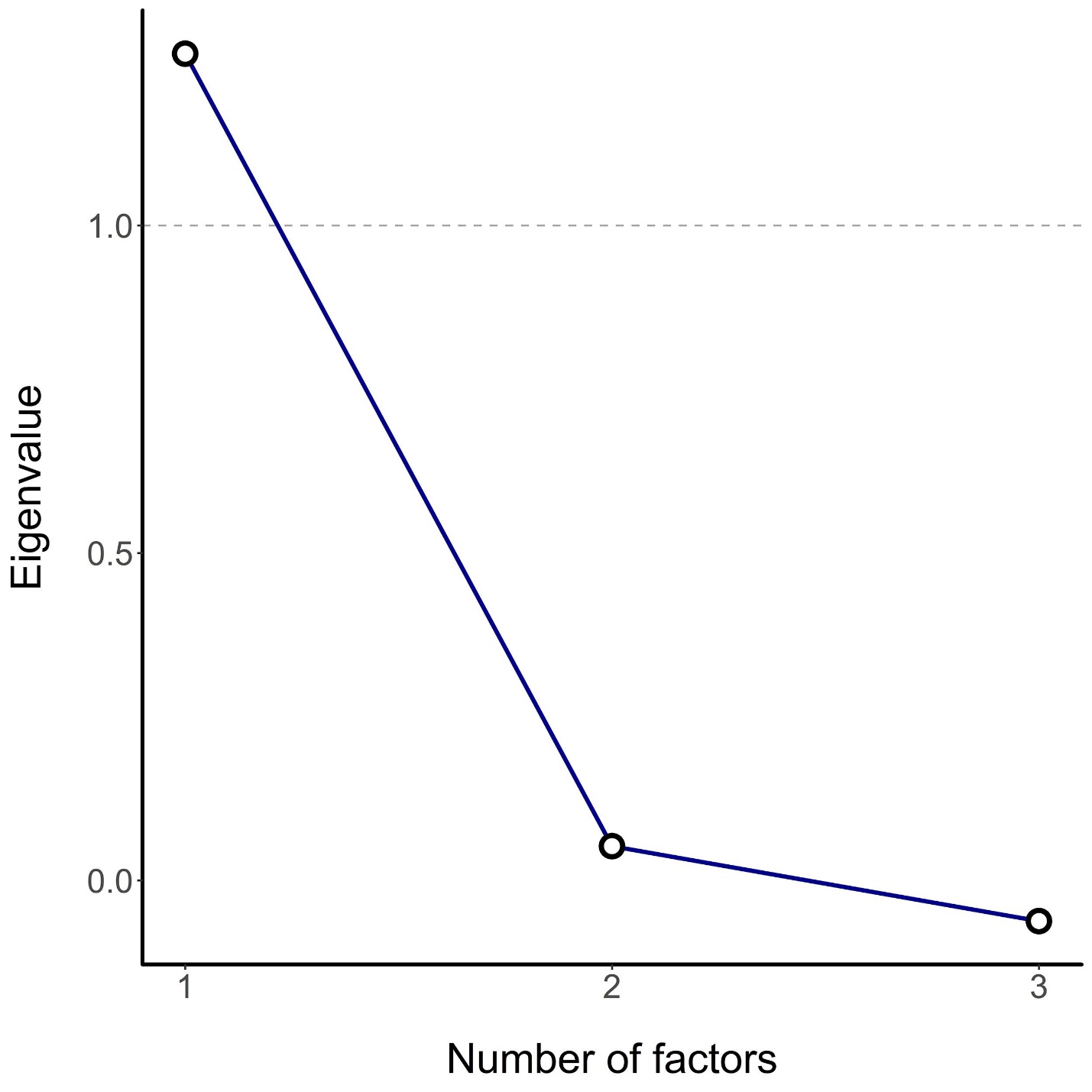

*Figure S1.* *Scree plot for factor analysis of pre-stimulation baseline dopamine measures. The one factor solution yielded the largest eigenvalues indicating it was the best fit to the data.*
